# Influenza A virus superinfection potential is regulated by viral genomic heterogeneity

**DOI:** 10.1101/286070

**Authors:** Jiayi Sun, Christopher B. Brooke

## Abstract

Defining the specific factors that govern the evolution and transmission of influenza A virus (IAV) populations is of critical importance for designing more effective prediction and control strategies. Superinfection, the sequential infection of a single cell by two or more virions, plays an important role in determining the replicative and evolutionary potential of IAV populations. The prevalence of superinfection during natural infection, and the specific mechanisms that regulate it, remain poorly understood. Here, we used a novel single virion infection approach to directly assess the effects of individual IAV genes on superinfection efficiency. Rather than implicating a specific viral gene, this approach revealed that superinfection susceptibility is determined by the total number of viral genes expressed, independent of their identity. IAV particles that expressed a complete set of viral genes potently inhibit superinfection, while semi-infectious particles (SIPs) that express incomplete subsets of viral genes do not. As a result, virus populations that contain more SIPs undergo more frequent superinfection. These findings identify both a major determinant of IAV superinfection potential and a prominent role for SIPs in promoting viral co-infection.

## Introduction

Influenza A viruses (IAV) are estimated to cause hundreds of thousands of deaths across the world every year during seasonal epidemics, despite widespread pre-exposure and vaccination(1). In addition to the yearly burden of seasonal influenza viruses, novel zoonotic IAV strains periodically emerge into humans from swine or birds, triggering unpredictable pandemics that can dramatically increase infection and mortality rates (2). Defining the specific factors that influence the evolution of influenza viruses is critical for designing more effective vaccines, therapeutics, and surveillance strategies.

The prevalence of co-infection can play an enormous role in determining the replicative and evolutionary potential of IAV populations. This is a function both of the segmented nature of the viral genome and the enormous amount of genomic heterogeneity present within IAV populations(3,4). Co-infection allows reassortment, the production of novel viral genotypes through the intermixing of the individual IAV genome segments (5,6). Reassortment events have contributed to the emergence of every major influenza pandemic of the past century(7). Co-infection also facilitates the complementation and productive replication of the semi-infectious particles (SIPs) that make up the majority of IAV populations (8–12). Finally, increasing the frequency of co-infection can accelerate viral replication kinetics and virus output by increasing the average multiplicity of infection (MOI)(13–15). Thus to better understand how IAV populations transmit and evolve, we must identify the specific host and viral factors that govern co-infection.

One of the primary means by which co-infection can occur is superinfection, the sequential infection of one cell by multiple viral particles. For some viruses, superinfection appears to occur freely(16,17). In contrast, a diverse range of viruses actively inhibit superinfection through a variety of mechanisms, a phenomenon known as superinfection exclusion (SIE)(18–26). The only in-depth study to date of IAV superinfection concluded that the viral neuraminidase (NA) protein acts to potently and rapidly inhibit IAV superinfection by depleting infected cells of the sialic acid receptors required for viral entry (27). More recently, Dou et al. reported a narrow time window during which IAV superinfection was possible(13). The existence of a potent mechanism of IAV SIE is at odds with both the frequent co-infection observed in a variety of experimental settings, and the widespread occurrence of reassortment at the global scale(28–33). Marshall et al. showed that superinfection up to 8 hours after primary infection leads to robust coinfection and reassortment in cell culture(34). Widespread co-infection and complementation have also been observed in the respiratory tracts of IAV-infected mice and guinea pigs(9,35). Collectively, these results suggest that IAV superinfection can be restricted, but to what extent and through which specific mechanisms remains a crucial open question.

Here, we reveal that IAV superinfection potential is directly regulated by the extent of genomic heterogeneity within the viral population. We observed that superinfection susceptibility is determined in a dose-dependent fashion by the number of viral genes expressed by the initially infecting virion, regardless of their specific identity. Further, we show that superinfection occurs more frequently in IAV populations with more SIPs compared with those with fewer. Finally, we demonstrate that SIE is mediated by the presence of active viral replication complexes, and is completely independent of gene coding sequence. Altogether, our results reveal how genomic heterogeneity influences IAV superinfection potential, and demonstrate how SIPs can modulate collective interactions within viral populations.

## Results

### Influenza virus SIE occurs in multiple cell types and is independent of type I interferon secretion

A previous study of IAV SIE concluded that NA expression completely blocks susceptibility to superinfection by 6 hours post-infection (hpi)(27). To explore the potential mechanisms of IAV SIE in greater detail, we developed a flow cytometry-based assay that allows us to precisely measure the effects of previous infection on superinfection efficiency. To clearly identify cells infected by the first virus, the superinfecting virus, or both, we used two recombinant viruses that express antigenically distinct hemagglutinin (HA), NA, and NS1 proteins that we could distinguish using specific monoclonal antibodies (mAbs) that we had on hand (**Fig S1**). For the primary infection, we used a recombinant version of the H1N1 strain A/Puerto Rico/8/34 (rPR8). For the secondary infection, we used a recombinant virus (rH3N2) that contained the HA and NA gene segments from the H3N2 strain A/Udorn/72, the NS gene segment from A/California/04/09, and the remaining 5 segments from PR8.

We first asked whether prior infection with rPR8 affected cellular susceptibility to superinfection with rH3N2. We infected MDCK cells with rPR8 at an MOI of <0.3 TCID50/cell, and at 3 hpi (all times post infection will be relative to the first virus added) we added the PR8-HA-specific neutralizing mAb H17-L2 to block secondary spread of rPR8 within the culture. At 6 hpi, we infected with rH3N2 at an MOI of <0.3 TCID50/cell. To prevent spread of both rPR8 and rH3N2, we added 20 mM NH4Cl at 9 hpi. In parallel, we performed simultaneous co-infections (0 hr) with rPR8 and rH3N2 to measure coinfection frequencies when SIE should not be possible. At 19 hpi we harvested cells and examined primary and secondary virus infection by flow cytometry, using H1 and H3 expression as markers of rPR8 and rH3N2 infection, respectively. We observed that the H3+ frequency within H1+ cells was significantly reduced when rPR8 infection preceded rH3N2 by 6 hrs compared with when rPR8 and rH3N2 were added simultaneously (**Fig 1A**). This indicated that rPR8 infection significantly reduces the susceptibility of cells to superinfection by 6 hpi.

**Figure 1.**
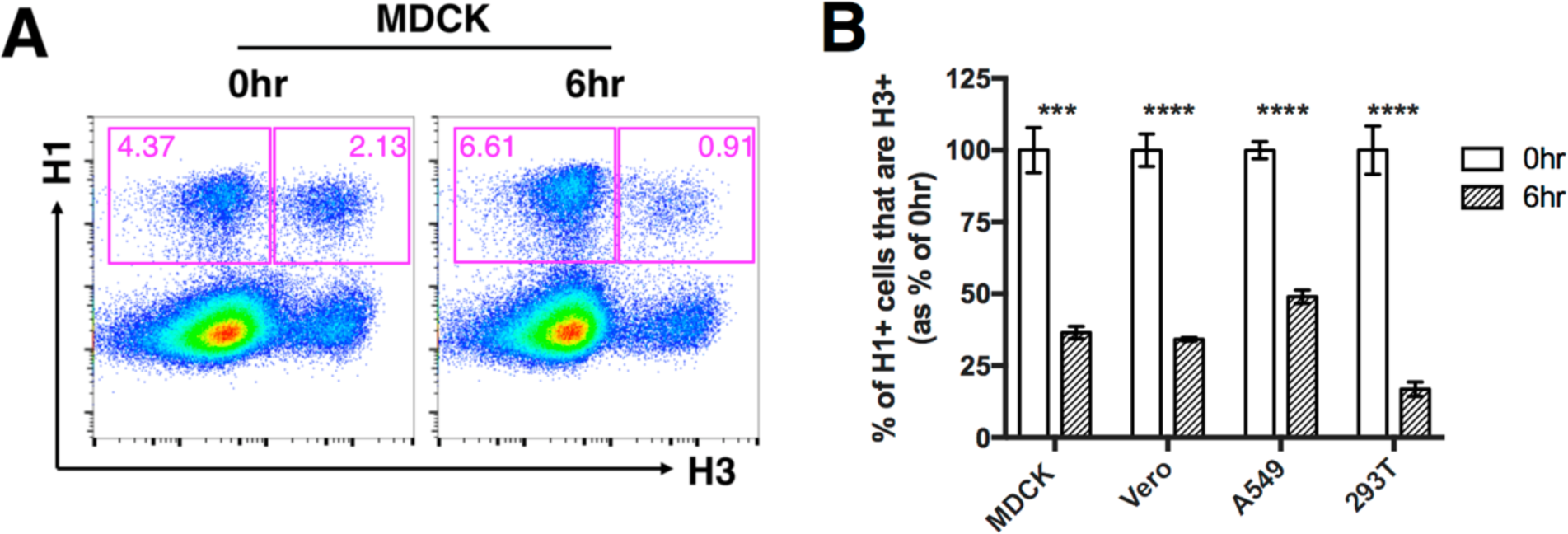
A 6-hr delay between primary infection and superinfection allows robust superinfection exclusion. The indicated mammalian cell lines were either simultaneously (0hr) or sequentially (6hr) infected with rPR8 and rH3N2 at MOI<0.3 TCID50/cell. **(A)** Representative FACS plots showing expression of H1 versus H3 within MDCK cells. **(B)** H3+ frequencies within H1+ cells following simultaneous or sequential infection, in the indicated cell lines. Values are calculated as the percentage of the mean 0hr value, and are presented as the mean (n=3 cell culture wells) ± standard deviations. \*\*\**P*<0.001; \*\*\*\**P*<0.0001; *t* test.

We next asked whether the SIE effect was cell type specific, and whether it depended upon activation of the type I interferon (IFN) system. We performed the same experiment as above in MDCK cells, A549 cells, 293T, and Vero cells (which are incapable of type I IFN secretion)(36,37). We observed that the extent of SIE was comparable between all cell lines tested, suggesting that SIE occurs in multiple distinct cell types, and does not depend upon IFN secretion (**Fig 1B; Fig S2**).

### Viral neuraminidase expression does not fully explain the SIE phenotype

In an attempt to confirm the previously reported role for NA activity in SIE, we directly measured the effect of NA expression on IAV SIE in our system(27). We took advantage of our previous observation that IAV populations consist primarily of SIPs that fail to express one or more viral genes(8). When carrying out the primary infection at low MOI, we generate populations of infected cells that are either positive or negative for expression of a given viral gene. We can then assess the effects of specific viral proteins on superinfection susceptibility by comparing superinfection frequencies between infected cells that do or do not express the protein in question.

We performed the same superinfection experiment as described above in MDCK cells, and at 19 hpi, harvested and stained with mAbs against H1, N1, and H3. To compare rPR8 infected cells that did or did not express NA, we individually gated cells into H1+N1+ and H1+N1-subpopulations (**Fig 2A**). Comparison of H3+ frequencies between H1+N1+ and H1+N1-cells revealed that NA expression was clearly associated with decreased susceptibility to superinfection in rPR8-infected cells (**Fig 2B**). This finding was consistent with the previously reported role for NA in IAV superinfection exclusion(27). Importantly, while SIE was most pronounced in the H1+N1+ cells, we also observed a significant decrease in superinfection susceptibility within the H1+N1-cell population by 6hpi, suggesting that viral factors other than NA also act to restrict superinfection.

**Figure 2.**
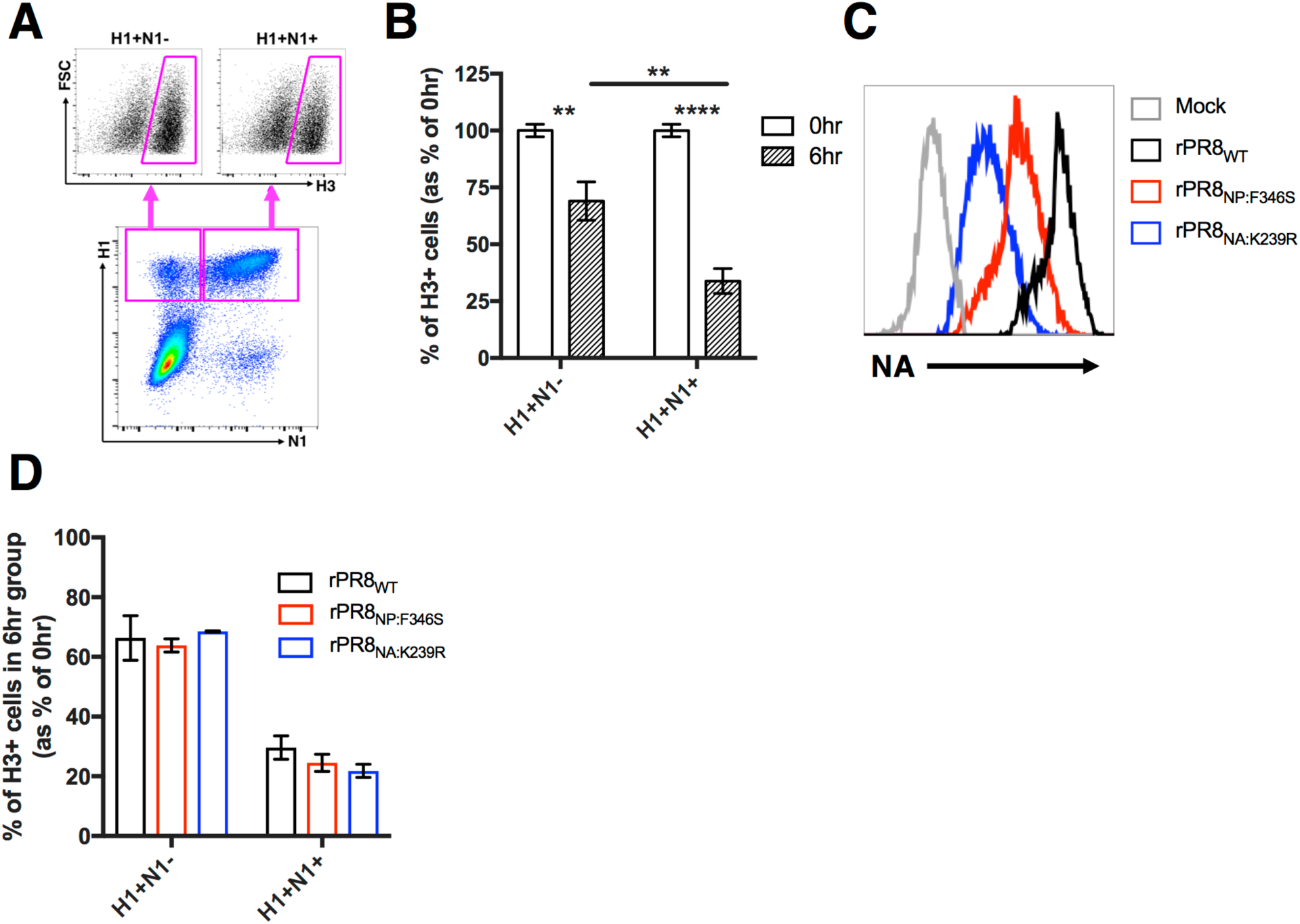
Superinfection is more potently inhibited in NA positive cells, independent of NA expression level. MDCK cells were infected with rPR8_WT_, rPR8_NP:F346S_ or rPR8_NA:K239R_, and simultaneously (0hr) or sequentially (6hr) infected with rH3N2; all infections at MOI<0.3 TCID50/cell. **(A)** Representative FACS plots comparing H3+ frequencies between H1+N1-and H1+N1+ cells. **(B)** H3+ frequencies within H1+N1-and H1+N1+ cells following simultaneous (0hr) or sequential (6hr) infection. Values are calculated as the percentage of the mean 0hr value. **(C)** Comparison of NA expression within H1+N1+ cells between rPR8_WT_, rPR8_NP:F346S_ and PR8_NA:K239R_ at 19 hpi. **(D)** Comparison of H3+ frequencies within H1+N1-and H1+N1+ cells following sequential infection (6hr) between the three indicated viruses. Values are normalized to H3+ frequencies in simultaneous infection (0hr) controls. For data in Fig B and D, mean values (n=3 cell culture wells) ± standard deviations are shown. \*\**P*<0.01; \*\*\*\**P*<0.0001; *t* test.

Relative NA activity can vary significantly between IAV strains(38). If NA activity inhibits IAV superinfection, we hypothesized viruses that express less NA would undergo more frequent superinfection. To define the quantitative relationship between NA expression and SIE, we examined the effects of two substitutions (NP:F346S and NA:K239R) that decrease cellular NA expression relative to wild-type PR8 on superinfection efficiency(9,39) (**Fig 2C**). Surprisingly, these mutants did not exhibit higher superinfection frequencies than wild-type PR8 (**Fig 2D**). These results suggest that SIE can be mediated by NA gene expression, but is not significantly influenced by relative NA expression levels.

### Superinfection susceptibility is determined by the number of viral genes expressed, not their identity

Based on our observation that superinfection was also inhibited within H1+N1-cells (**Fig 2B**), we hypothesized that expression of other viral gene products can also inhibit superinfection. We examined the effects of HA and NS1 expression on superinfection susceptibility, using rPR8-specific mAbs. Surprisingly, we found that both HA and NS1 expression within rPR8-infected cells were also associated with significant decreases in superinfection by rH3N2, comparable to the effect associated with NA expression (**Fig 3A**,**B**). To further dissect the effects of viral gene expression patterns on SIE, we individually gated all seven possible combinations of HA, NA, and NS1 expression by rPR8 (HA+NA+NS1+, HA+NA+, HA+NS1+, NA+NS1+, HA+, NA+, NS1+) and directly compared rH3N2 infection frequencies between them (gating scheme: **Fig. S3**). We observed that the fraction of cells superinfected with rH3N2 was inversely correlated with the number of rPR8 genes expressed, regardless of their specific identities (**Fig 3C**,**D**). Thus, susceptibility to IAV superinfection is determined by the number of viral genes expressed in the host cell, rather than their specific identity.

**Figure 3.**
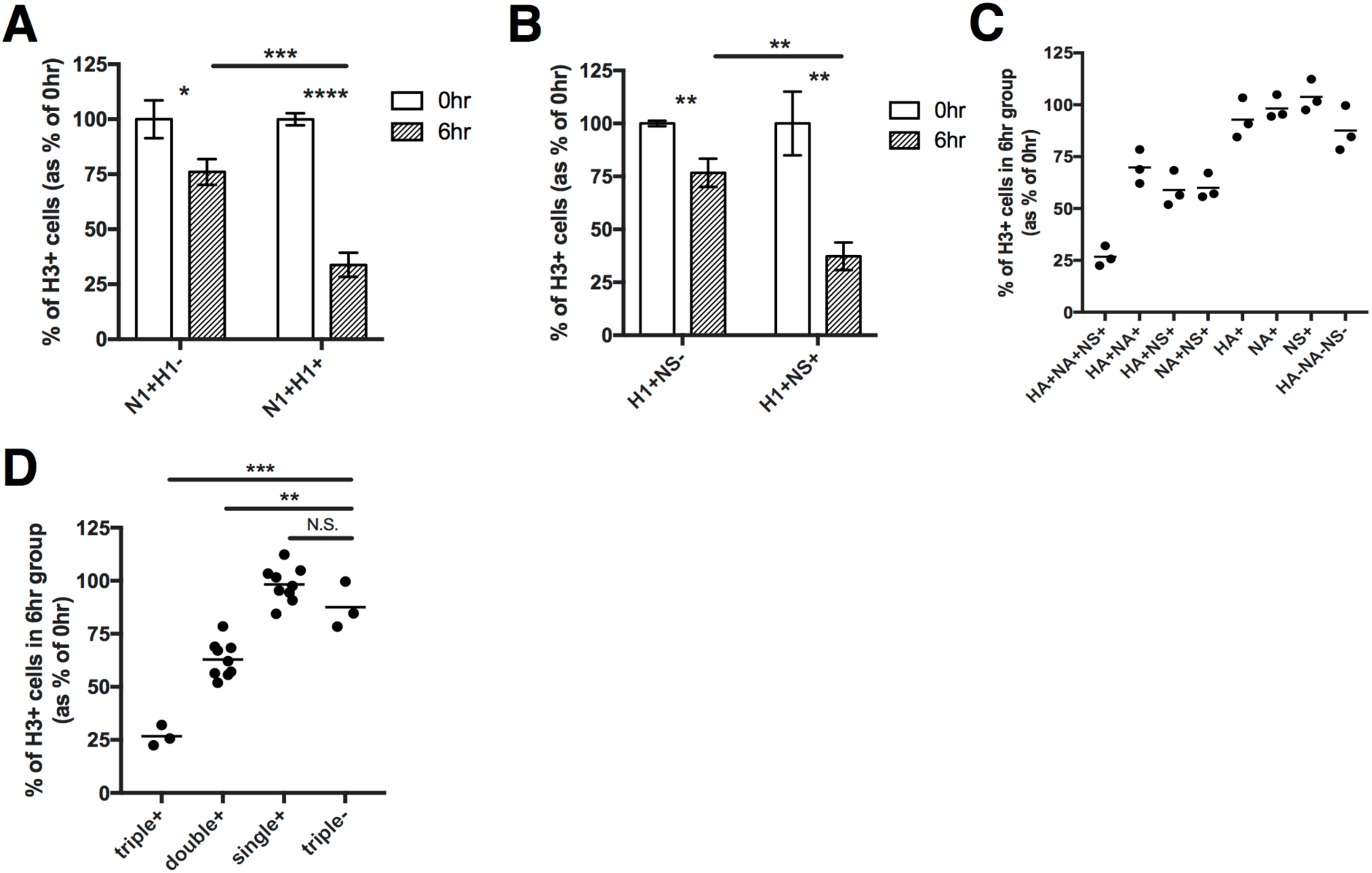
Superinfection is more frequent in cells that express fewer viral genes. MDCK cells were infected with rPR8, and simultaneously (0hr) or sequentially (6hr) infected with rH3N2; all infections at MOI<0.3 TCID50/cell. **(A)** H3+ frequencies within N1+H1-and N1+H1+ cells following simultaneous (0hr) or sequential (6hr) infection. Values are calculated as the percentage of the mean 0hr value. **(B)** H3+ frequencies within H1+NS1-and H1+NS1+ cells following simultaneous (0hr) or sequential (6hr) infection. Values are calculated as the percentage of the mean 0hr value. **(C)** Comparison of H3+ percentages between 8 cell populations gated based on the expression of the indicated combinations of 3 PR8 gene products (HA, NA, NS1). Data represents the values obtained from 6hr samples normalized to the values obtained from 0hr control samples. Each data point represents the value for the indicated cell population within a single cell culture well. **(D)** Data in C grouped by total numbers of expressed viral gene products rather than their specific identity. \*\**P*<0.01; \*\*\**P*<0.001; *t* test.

### Superinfection is more prevalent in IAV populations with more SIPs

If the number of viral genes expressed in a cell determines superinfection susceptibility, then decreasing the average number of functional viral genes successfully delivered by individual virions should increase the overall incidence of superinfection. We tested this by artificially decreasing the functional gene segment content of rPR8 through exposure to UV irradiation(40). Exposure to low dose UV irradiation generates SIPs that carry gene-lethal UV-induced lesions at frequencies proportional to genome segment length. Based on our previous findings, we hypothesized that superinfection frequencies would increase with longer exposure of rPR8 to UV.

We UV irradiated (302nm) rPR8 for either 30s or 60s, and confirmed that the TCID50 concentration was reduced and the SIP concentration was increased as a function of treatment duration (**Fig 4A-C**). We then performed superinfection assays as before, comparing rH3N2 superinfection frequencies between untreated and UV-irradiated rPR8 in MDCK cells. To fairly compare superinfection frequency between viral populations with differing particle-to-infectivity ratios, we normalized our rPR8 infections based on equivalent numbers of particles capable of expressing NA (NA-expressing units; NAEU)(9).

**Figure 4.**
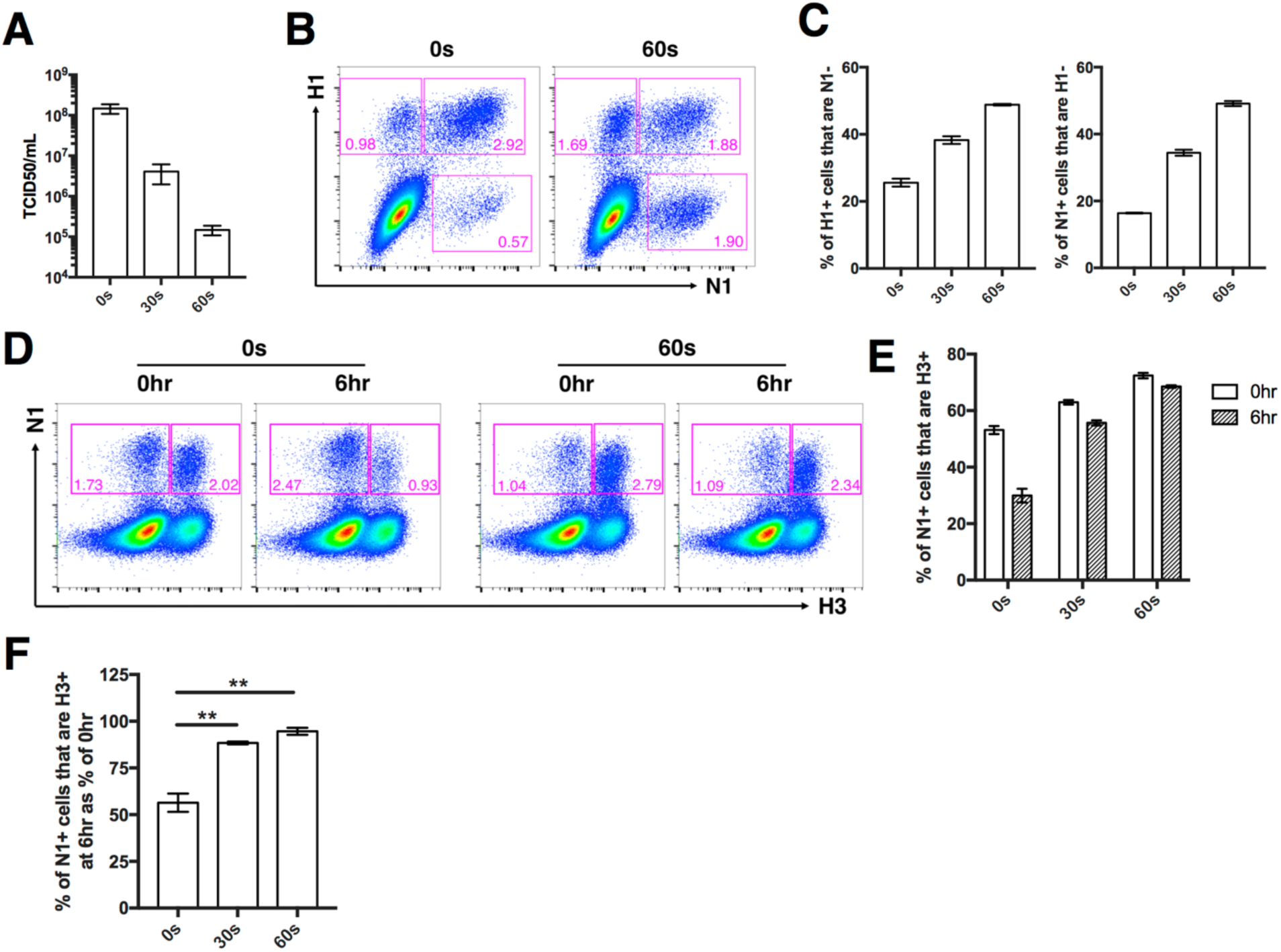
Superinfection is more common in viral populations that contain more SIPs. rPR8 was irradiated with 302nm UV lamp for 30s or 60s. **(A)** TCID50 titers of untreated (0s), 30s, and 60s UV-treated rPR8. Data from two independent experiments are shown. **(B)** Representative FACS plots showing HA and NA expression patterns of untreated (0s) and 60s UV-treated rPR8. **(C)** Quantification of HA/NA co-expression in infected cells from (B). **(D-F)** MDCK cells were infected with untreated (0s), 30s, or 60s UV-treated rPR8 at an MOI of 0.04 NAEU/cell, and simultaneously (0hr) or sequentially (6hr) infected with rH3N2 at MOI<0.3 TCID50/cell. **(D)** Representative FACS plots showing expression of N1 versus H3 in cells infected with either untreated (0s) or 60s UV-treated rPR8. **(E)** rH3N2 infection percentages within N1+ cells infected with either untreated (0s), 30s, or 60s UV-treated rPR8. **(F)** Data from 6hr samples in E shown as % of 0hr control samples. For Fig C, E, and F, mean values (n=3 cell culture wells) ± standard deviations are shown. \*\**P*<0.01; *t* test.

We first examined the effect of UV treatment on superinfection when rPR8 and rH3N2 were added to cells simultaneously (0hr). This was a critical control because UV treatment can increase the measured incidence of co-infection independent of SIE effects, purely by creating a larger pool of SIPs that only show up in our assays when complemented by secondary infection(40). Consistent with this, we observed a small increase in co-infection frequency with UV treatment when both viruses were added simultaneously (**Fig 4D**,**E**). When rH3N2 was added 6 hours after rPR8 however, we observed a much more pronounced increase in superinfection frequency with UV treatment, consistent with our hypothesis that superinfection is regulated by the proportion of SIPs present within the viral population (**Fig 4D-F**).

### SIE is mediated by active IAV replication complexes, and is independent of gene coding sequence

Our data reveal that IAV superinfection potential is determined by the number of viral genes expressed within a cell, independent of their specific identity. This suggests that the viral gene products themselves are dispensible for SIE. We thus hypothesized that active replication and/or transcription of viral RNAs by the viral replicase complex is responsible for decreasing cellular susceptibility to subsequent infection. To test this, we co-transfected 293T cells with pDZ vectors encoding the individual viral replicase proteins (PB2, PB1, PA, and NP) together with a pHH21 vector encoding either the HA vRNA gene segment (HA_vRNA_) or a vRNA-derived reporter gene segment in which the eGFP ORF is flanked by the 5’ and 3’ UTR sequences from the NA segment (eGFP_vRNA_). These UTR sequences are required for replication and transcription of the reporter RNA by the viral replicase. 24 hours post transfection, we infected cells with rH3N2 at an MOI of 0.2 TCID50/cell and measured infectivity at 8 hpi using an M2-specific mAb.

Infection frequencies were decreased ~50% in cells expressing the replicase components plus the eGFP_vRNA_ construct, compared with control cells transfected with the replicase-expressing constructs plus an empty pHH21 vector (**Fig 5A**). When comparing rH3N2 infectivity between co-transfected cells (eGFP+, HA+) and un-co-transfected cells (eGFP-, HA-) within the same culture wells, the inhibitory effects mediated by eGFP_vRNA_ or HA_vRNA_ expression were comparable (**Fig 5B**,**S4**). Importantly, this effect was not seen when we left out the plasmid encoding PA (RNP_PA-_) or used an eGFP reporter RNA that lacked the viral UTR sequences (eGFP_ORF_) (**Fig 5A**). Altogether, these data indicate that the inhibition of infection requires both an intact replicase complex and an RNA template containing the viral UTR sequences, but not the viral coding sequence.

**Figure 5.**
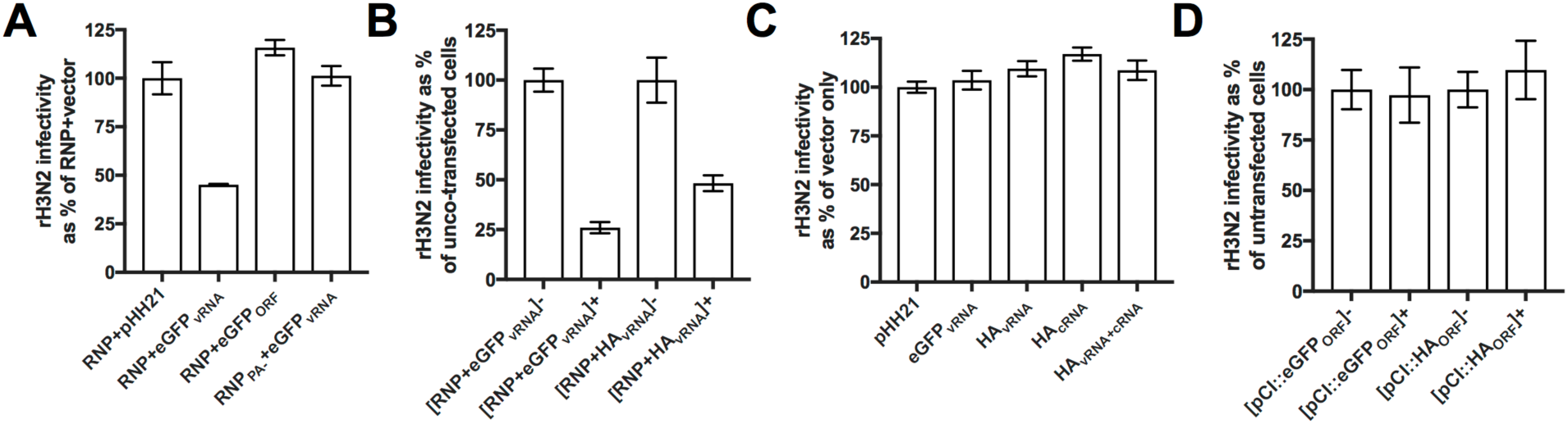
Activity of viral replication complexes is responsible for the inhibition on subsequent infection, not those viral gene products. **(A)** 293T cells were co-transfected with plasmids encoding the viral replicase complex proteins PB2, PB1, PA, and NP (RNP), together with 1 μg of either empty pHH21 vector, pHH21::eGFP_vRNA_ (eGFP ORF flanked with NA UTRs), or pHH21::eGFP_ORF_ (eGFP ORF with no UTR sequence). Control cells were co-transfected with plasmids encoding the viral replicase complex minus PA, which was replaced by empty vector (RNP_PA-_), together with 1 μg of pHH21::eGFP_vRNA_. 24 hours post-transfection, cells were infected with rH3N2 at MOI=0.2 TCID50/cell. At 8 hpi, cells were harvested and assessed for rH3N2 infectivity via flow cytometry. rH3N2 infectivity are normalized to RNP+vector. **(B)** Similar experiment in (A) with co-transfection of RNP plasmids plus 1 μg of pHH21::eGFP_vRNA_ or pHH21::HA_vRNA_. rH3N2 infectivity in co-transfected cells (eGFP+, HA+) are normalized to un-co-transfected cells (eGFP-, HA-) within the same culture wells. **(C)** Similar experiment in (A) with transfection of empty pHH21 vector, pHH21::eGFP_vRNA_, pHH21::HA_vRNA_, pHH21::HA_cRNA_, or pHH21::HA_vRNA_ plus pHH21::HA_cRNA_. rH3N2 infectivity are normalized to empty pHH21 vector. **(D)** Similar experiment in (A) with transfection of pCI vector, pCI::eGFP_ORF_ or pCI::HA_ORF_. rH3N2 infectivity in transfected cells (eGFP+, HA+) are normalized to un-transfected cells (eGFP-, HA-) within the same culture wells. Mean values (n=2 cell culture wells) ± standard deviations are shown.

Our data demonstrate that IAV SIE is mediated by the specific activity of viral replication complexes. One potential explanation is that large amounts of recently-synthesized negative sense vRNA within the cell might outcompete incoming genome segments for replication and expression. To test this, we transfected 293T cells with a pHH21 vector that overexpresses the eGFP_vRNA_ segment, and measured susceptibility to rH3N2 infection 24 hours later using an NP-specific mAb. Compared to the empty vector control, we observed no effect of eGFP_vRNA_ vRNA overexpression on cellular susceptibility to infection (**Fig 5C**). Similarly, we observed no effect when we overexpressed the cRNA or vRNA forms of the HA gene segment, either individually or together (**Fig 5C**). Another potential explanation is that viral mRNA or protein overexpression might inhibit subsequent infection. To test this, we transfected 293T cells with pCI vectors that overexpress mRNA and protein of eGFP and HA, and measured susceptibility to rH3N2 infection 24 hours post transfection using the M2-specific mAb. Compared to empty vector control, mRNA/protein overexpression of eGFP or HA had no effect on the following infection (**Fig 5D S5A, S5B**). Altogether, these data demonstrate that IAV SIE is driven by the presence of active viral replication complexes, rather than the protein or nucleic acid products of viral genes processed by those complexes.

## Discussion

Superinfection plays an enormous role in influencing the outcome of IAV infection, both by promoting reassortment and by facilitating the multiplicity reactivation of SIPs and defective interferring particles (4). Despite this importance, the specific factors that govern the occurrence of superinfection have remained obscure. Here, we reveal that IAV superinfection susceptibility is directly regulated by the number of viral genes expressed by a virion, regardless of their specific identity or function. This effect depends upon the presence of active viral replication complexes, but not their nucleic acid or protein products. Critically, we demonstrate that the presence of SIPs within viral populations significantly increases the frequency of superinfection. This represents a completely novel mechanism of viral superinfection exclusion and identifies a clear mechanistic consequence of the enormous genomic heterogeneity within IAV populations.

The only other published study that examines IAV SIE in detail concluded that NA expression mediates SIE by depleting the pool of available sialic acid receptors on the cell surface(27). In this study, we directly quantified the contribution of NA expression to SIE during IAV infection, and found that the SIE effect of NA expression is actually comparable to that of other viral genes and that overall genomic content is the primary determinant of superinfection potential. The conclusions of the Huang et al. study were based primarily on two observations: (*1*) overexpression of NA within cells rendered them refractory to infection by an HA-pseudotyped virus, and (*2*), IAV superinfection only occured when cells were treated with NA inhibitors (NAIs)(27). While we cannot conclusively explain the discrepancies between the two studies, we can offer a couple of plausible explanations. First, the cellular overexpression studies in Huang et al. likely involved levels of cellular NA expression that are far beyond those seen during IAV infection. In fact, we were also able to observe a similarly potent restriction of IAV infection following plasmid-driven NA overexpression (data not shown); however, this result did not reflect what we observed during viral infection (**Fig 2**). Second, the observation that NAI treatment dramatically increases superinfection frequencies may be explained by the effects of cell death. In their experiments, Huang et al. infected cells at a relatively high MOI, did not block secondary spread of the virus within cultures, and assessed superinfection frequency at 20 hpi or later. Under these conditions, many of the initially infected cells will be dead or dying and thus lost from the analysis. This may be especially true of superinfected cells, which will tend to be infected at a higher than average effective MOI. Even under low MOI conditions, we had to limit the timeframe of our experiments and block secondary spread of virus to prevent cell death from skewing our results. NAI treatment may act to help preserve co-infected cells so that they are detected at the endpoint of the experiment, thus increasing the measured superinfection rate.

Our results reveal that SIE is mediated by multiple IAV genes in a dose-dependent fashion. The surprising irrelevance of the specific IAV gene segments involved is explained by our finding that the viral coding sequence of a gene segment can be replaced with that of eGFP without any loss of inhibitory effect. This suggests a direct role for viral replicase activity itself in triggering SIE, rather than any effect of the viral gene segments themselves. The specific mechanism by which the activity of viral replicase complexes may inhibit subsequent infection remains unclear, however one potential explanation is that viral replication complexes trigger a dose-dependent intrinisic host anti-viral response. While our experiments in Vero cells demonstrate that the secretion of type I IFN is not required for SIE, they do not preclude the involvement of type I IFN-independent mechanisms. These could include either the type III IFN-mediated induction of anti-viral effectors, or the engagement of completely IFN-independent anti-viral mechanisms(41– 43). Future studies are aimed at delineating the role of the host in the regulation of IAV superinfection.

Our results demonstrate that SIPs can directly influence the prevalence of superinfection, and thus potentially the frequency of reassortment. Fonville et al. used a similar UV irradiation-based method as shown here to demonstrate that increasing the frequency of SIPs within a viral popuation increases the overall reassortment rate(40). The explanation given for this effect was that increasing the abundance of SIPs increases the proportion of the viral population that depends upon co-infection to replicate. As a result, within a certain MOI range, a greater share of productively infected cells will be co-infected and subject to reassortment. In our study, we confirmed this effect by observing a slight increase in co-infection frequency with increasing UV dose when rPR8 and rH3N2 were added simultaneously (**Fig 4E**). When we controlled for this however, we still observed a significant increase in superinfection frequency as we increased the proportion of SIPs through UV treatment (Fig 4D-F). Thus, the relationship between SIPs and SIE that we describe here is completely independent of the increased multiplicity reactivation observed by Fonville et al. and likely represents the effects of decreasing the strength of SIE. Between these two studies, it is clear that SIPs can modulate the frequency of IAV co-infection and reassortment though at least two distinct mechanisms.

IAV strains can differ significantly in the relative production and gene expression patterns of SIPs(8,9). This raises the possibility that strains with distinct SIP production phenotypes may differ in their reassortment potential, given the influence of SIPs over coinfection and reassortment frequencies. If this is the case, it would suggest a significant role for SIPs production in governing the evolutionary potential of IAV populations.

The relationship between viral gene expression patterns and superinfection exclusion that we report here demonstrates that viral genomic heterogenity has distinct functional consequences during infection. A crucial implication is that all infected cells cannot be thought of as equal, but may in fact exhibit distinct phenotypes based on the number and identity of viral genome segments they harbor. The relationship between viral genomic heterogeneity and the phenotypic diversity of infected cells likely extends to other cellular features beyond superinfection susceptibility.

It remains to be seen whether the relationship between viral gene dose and superinfection susceptibility that we describe here exists for other segmented viruses besides IAV. Beyond the segmented viruses, it has become increasingly clear that collective interactions mediated by cellular co-infection significantly influence the replicative and evolutionary dynamics of non-segmented viruses as well(44). More work is needed to better understand the factors that govern co-infection for different virus families, both *in vitro* and *in vivo*.

In summary, our work reveals a unique mechanism of IAV superinfection regulation that is governed by viral genomic heterogeneity. Critically, we show that the abundance of SIPs within a viral population directly influences the prevalence of superinfection; suggesting that IAV strains may differ in their superinfection potential, and thus potential for reassortment. This finding has significant consequences for understanding the evolutionary potential of different IAV genotypes with varying SIP phenotypes. More broadly, we demonstrate how genomic diversity within viral populations can have clear functional consequences during infection.

## Methods

### Plasmids

The A/Puerto Rico/8/34 and A/Udorn/72 reverse genetics plasmids were generous gifts from Dr. Adolfo Garcia-Sastre and Dr. Kanta Subbarao, respectively. The pCI vector wa s graciously provided by Dr. Joanna Shisler. The pHH21::eGFP_vRNA_ (eGFP ORF flanked by the NA UTRs) was kindly gifted by Dr. Andrew Mehle. Generation of pHH21::HA_vRNA_ has been previously described(9). The following primer pairs were used to generate the indicated constructs. pHH21::eGFP_ORF_ (BsmBI): 5’ CGTCTCCTATTTTACTTGTACAGC TCG; 3’ CGTCTCCGGGATGGTGAGCAAGGGC. pHH21::HA_cRNA_ (BsmBI): 5’CGTCTCA TATTAGCAAAAGCAGG; 3’ CGTCTCAGGGAGTAGAAACAAGGG. pCI::eGFP_ORF_ (Eco RI/SalI): 5’ AGAATTCATGGTGAGCAAGG; 3’AGTCGACTTACTTGTACAGC. pCI::HA_OR_ F (EcoRI/SalI): 5’ AGAATTCATGGAAGATTTTGTGCG; 3’ AGTCGACCTAACTCAATGC ATGTGT.

### Cells

Madin-Darby canine kidney (MDCK) and human embryonic kidney HEK293T (293T) cells were maintained in Gibco’s minimal essential medium with GlutaMax (Life Technologies). Vero cells were maintained in Dulbecco’s modified eagle medium (Life Technologies). Human lung epithelial A549 cells were maintained in Gibco’s F-12 medium (Life Technologies). MDCK, vero, and A549 cells were obtained from Dr. Jonathan Yewdell; 293T cells were obtained from Dr. Joanna Shisler. All media were supplemented with 8.3% fetal bovine serum (Seradigm). Cells were grown at 37°C and 5% CO2.

### Viruses

Recombinant A/Puerto Rico/8/1934 (rPR8) and rH3N2 viruses were generated using 8-plasmid rescue systems. The rH3N2 virus is a reassortant with the HA and NA segments from A/Udorn/72 (H3N2), the NS segment from A/California/04/09 (H1N1), and the other 5 segments from PR8. The rPR8 clones differ from the published sequence (GenBank accession nos. AF389115–AF389122) at two positions: PB1 A549C (K175N) and HA A651C (I207L) (numbering from initiating Met). Molecular clone-derived mutants (rPR8 NP:F346S and rPR8 NA:K239R) were generated using standard site-directed PCR mutagenesis. All viruses were rescued by transfecting sub-confluent 293T cells with 500ng of each of the appropriate reverse genetics plasmids using JetPRIME (Polyplus) according to the manufacturer’s instructions. Plaque isolates derived from rescue supernatants were amplified into seed stocks in MDCK cells. Working stocks were generated by infecting MDCK cells at an MOI of 0.0001 TCID50/cell with seed stock and collecting and clarifying supernatants at 48 hpi. All viral growth was carried out in MEM with 1 µg/mL trypsin treated with L-(tosylamido-2-phenyl) ethyl chloromethyl ketone (TPCK-treated trypsin; Worthington), 1mM HEPES, and 100 µg/mL gentamicin. Virus stocks were titered via standard tissue culture infectious dose 50 (TCID50) assay.

### Superinfection assay

For the 6hr sequential infection group, confluent mammalian cells (MDCK, vero, A549, or 293T) in six-well plates were infected with rPR8 at MOI<0.3 TCID50/cell for 1 hour. 1 hour post-adsorption, monolayers were washed with PBS and incubated in serum-containing medium. At 3 hpi, neutralizing anti-PR8-HA mouse mAb H17-L2 (5 µg/mL) was added to cultures to prevent spread of rPR8. At 6 hpi, monolayers were superinfected with rH3N2 at MOI<0.3 TCID50/cell in the presence of H17-L2 (which does not interfere with rH3N2 infection, Fig S2). 1 hour post-adsorption, monolayers were washed with PBS and incubated in serum-containing medium with H17-L2. At 9 hpi of rPR8 (3 hpi of rH3N2), the media was changed to MEM with 50 mM Hepes and 20 mM NH4Cl to block spread of both viruses. At 19 hpi of rPR8 (13 hpi of rH3N2), monolayers were trypsinized into single-cell suspensions.

For the 0hr simultaneous infection group, cells were infected with a mixture of rPR8 and rH3N2 at the same MOIs as in 6hr superinfection group. At 3 hpi, the NH4Cl media was added to block viral spread and cells were harvested at 19 hpi.

All cells were simultaneously fixed and permeabilized using foxP3 fix/perm buffer (eBioscience). Fixed cells were stained with Alexa Fluor 488-conjugated mouse anti-H1 mAb H36-26 (which does not compete with H17-L2), Pacific Orange-conjugated mouse anti-N1 mAb NA2-1C1, Pacific Blue-conjugated mouse anti-NS1 mAb NS1-1A7 and Alexa Fluor 647-conjugated mouse anti-H3 mAb H14-A2 (All mAbs gifts of Dr. Jon Yewdell). After staining, cells were washed, run on a BD LSR II, and analyzed using FlowJo version 10.1 (Tree Star, Inc.).

### Quantification of NA expression

H1+N1+ MDCK cells infected with rPR8WT, rPR8 NP:F346S, or rPR8 NA:K239R in the superinfection assay (6hr group) were gated and histograms for NA expression were plotted. The geometric mean fluorescence intensities (GMFI) for NA were determined using FlowJo version 10.1 (Tree Star, Inc.).

### UV treatment and analysis

rPR8 stocks were placed in six-well plates on ice (500 ul/well). Plates were placed 5 cm underneath a 302nm UVP-57 handheld UV lamp (UVP) and irradiated for 30s or 60s. TCID50 titers and single virion expression patterns of untreated and UV-treated virus were determined on MDCK cells, and superinfection assays described above were performed using these viruses and rH3N2.

### Transfection assay

80% confluent 293T cells in six-well plates were transfected with the following plasmids using jetPRIME (Polyplus): RNP (500 ng each of pDZ::PB2, pDZ::PB1, pDZ::PA, and pDZ::NP) plus 1 μg of pHH21 vector; RNP plus 1 μg of pHH21::eGFP_vRNA_ (eGFP_ORF_ flanked with NA UTRs); RNP plus 1 μg of pHH21::eGFP_ORF_; RNP plus 1 μg of pHH21::HA_vRNA_; RNP_PA-_ (500 ng each of pDZ::PB2, pDZ::PB1, pDZ, and pDZ::NP) plus 1 μg of pHH21::eGFP_vRNA_; RNP_PA-_ plus 1 μg of pHH21::HA_vRNA_; 6 µg of pHH21 vector; 3 µg of pHH21 vector plus 3 µg of pHH21::eGFP_vRNA_; 3 µg of pHH21 vector plus 3 µg of pHH21::HA_vRNA_; 3 µg of pHH21 vector plus 3 µg of pHH21::HA_cRNA_; 3 µg of pHH21::HA_vRNA_ plus 3 µg of pHH21::HA_cRNA_; 3 µg of the pCI vector; 3 µg of pCI::eGFP_ORF_; 3 µg of pCI::HA_ORF_. All plasmid-encoded viral sequences were derived from PR8. 24 hours post transfection, monolayers were infected with rH3N2 at an MOI of 0.2 TCID50/cell. At 8 hpi, cells transfected by RNP+pHH21 and pCI plasmids were harvested and stained with Alexa Fluor 488-conjugated mouse anti-H1 mAb H36-26 and Alexa Fluor 647-conjugated mouse anti-M2 mAb O19. Cells transfected with pHH21 plasmids were permeabilized, fixed and stained with Alexa Fluor 647-conjugated mouse anti-NP mAb HB-65. After staining, cells were washed, run on a BD LSR II, and virus infection frequencies, as measured by fractions of M2+ or NP+ cells, were quantified using FlowJo version 10.1 (Tree Star, Inc.).

### Statistical analysis

Unpaired, two-sided student’s t-tests were applied to the data shown in Fig 1B Fig 2B Fig 3A Fig 3B and Fig 4F. An unpaired, two-sided Welch’s t-test was applied to the data shown in Fig 3D. All statistical analyses were performed with GraphPad Prism 7.0a.

## Acknowledgements

We are greatful to the other members of the lab for helpful discussions and critical reading of the manuscript. This work has been generously supported by the National Institute of Allergy and Infectious Diseases of the National Institutes of Health under Award Number K22AI116588, as well as startup funds from the University of Illinois.

**Supporting information**

**Figure S1. Expression of HA, NA, and NS1 by rPR8 and rH3N2 can be differentiated using specific mAbs**. MDCK cells were infected with rPR8 or rH3N2 at an MOI<0.3 TCID50/cell. At 19 hpi, cells were harvested, fixed, permeabilized, stained against H1 (H36-26), N1 (NA2-1C1), NS1 (1A7), and H3 (H14-A2), and run on an LSR II flow cytometer. Expression of H1 versus N1, and NS1 versus H3 are shown in representative FACS plots.

**Figure S2. Superinfection is inhibited in multiple cell lines**. H1 versus H3 expression in Vero cells, A549 cells, and 293T cells observed in the experiments described in Fig 1 are shown in as representative FACS plots.

**Figure S3. Gating scheme for measuring superinfection frequencies in cells that express different rPR8 genes**. Cells from the experiment described in Fig 3C,D were assessed for expression of HA, NA and NS sequentially. The fractions of H3+ cells (indicative of superinfection rates) were quantified and compared between each of the eight cell populations with the indicated expression patterns.

**Figure S4. Cells co-transfected with plasmids encoding viral replicase and viral vRNA are less susceptible to subsequent infection**. Cells from the experiments described in Fig 5A,B are assessed for expression of M2 (indicative of rH3N2 infection) versus eGFP and HA (indicative of co-transfection) in representative FACS plots.

**Figure S5. Overexpression of mRNA and protein does not inhibit subsequent infection. (A)** Cells from the experiment described in Fig 5D are assessed for expression of M2 (indicative of rH3N2 infection) versus eGFP and HA (indicative of transfection) in representative FACS plots. **(B)** rH3N2 infectivity in A are shown as % of pCI vector. Mean values (n=2 cell culture wells) ± standard deviations are shown.

