## Supplementary Materials for "Influenza A virus superinfection potential is regulated by viral genomic heterogeneity"

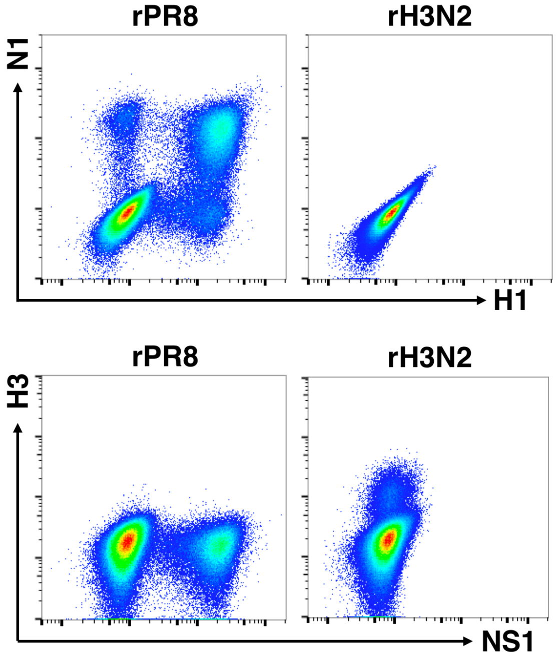


**Figure S1. Expression of HA, NA, and NS1 by rPR8 and rH3N2 can be differentiated using specific mAbs.** MDCK cells were infected with rPR8 or rH3N2 at MOI<0.3 TCID50/cell. At 19 hpi, cells were harvested, fixed, permeabilized, stained against H1, N1, NS1, and H3, and run on an LSR II flow cytometer. Expression of H1 versus N1, and NS1 versus H3 are shown in representative FACS plots.

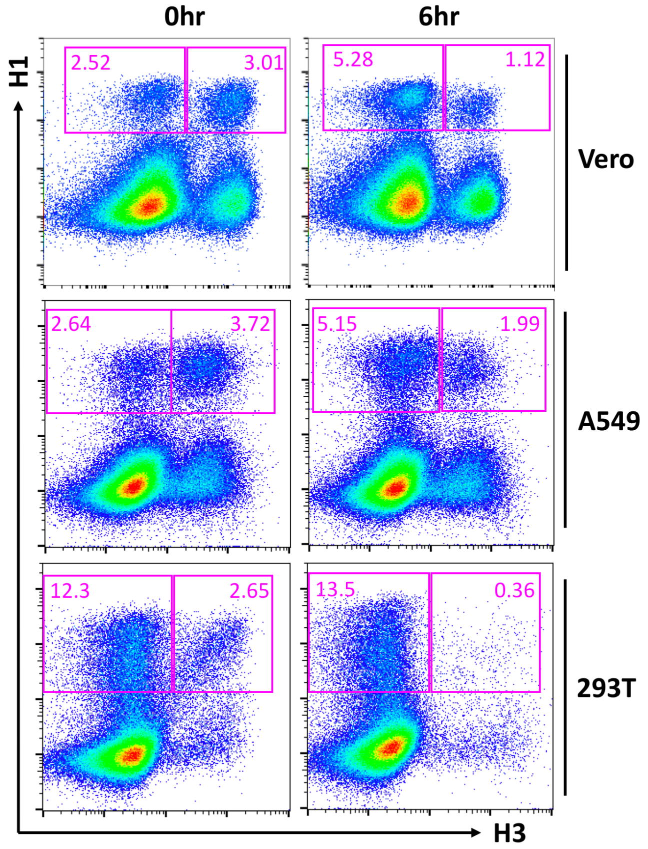


**Figure S2. Superinfection is inhibited in multiple cell lines.** H1 versus H3 expression in Vero cells, A549 cells, and 293T cells from the experiment described in Fig 1 are shown in representative FACS plots.

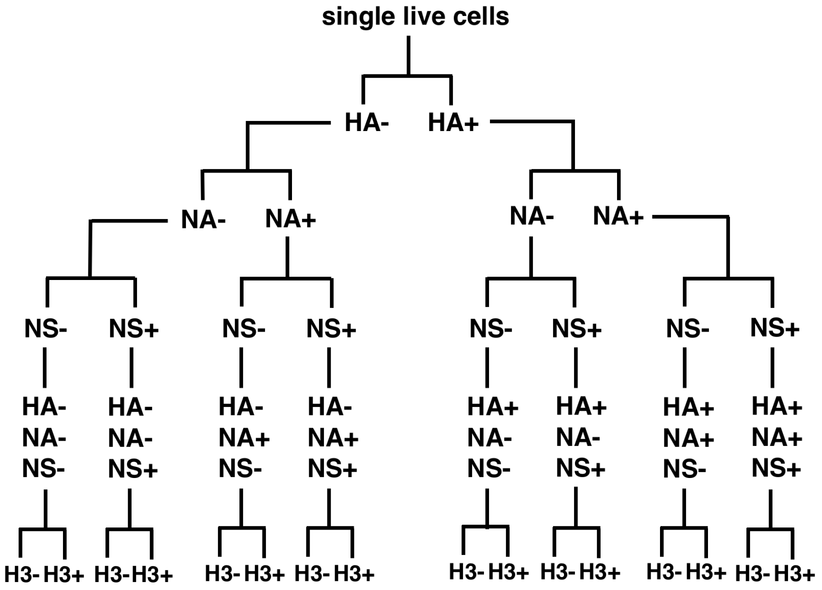


**Figure S3. Gating scheme for measuring superinfection frequencies in cells that express different rPR8 genes.** Cells from the experiment described in Fig 3C,D were assessed for expression of HA, NA, and NS sequentially. The fractions of H3+ cells (indicative of superinfection rates) were quantified and compared between each of the eight cell populations with the indicated expression patterns.

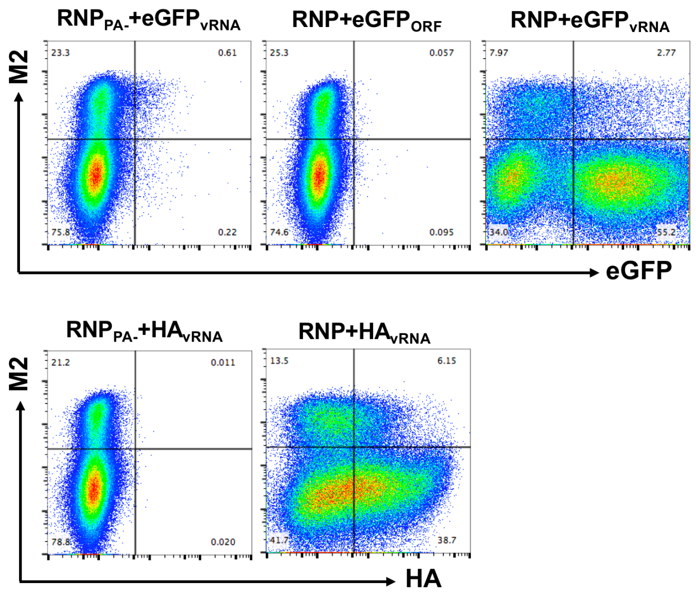


**Figure S4. Cells co-transfected with plasmids encoding viral replicase and viral vRNA are less susceptible to subsequent infection.** Cells from the experiments described in Fig 5A,B are assessed for expression of M2 (indicative of rH3N2 infection) versus eGFP and HA (indicative of co-transfection) in representative FACS plots.

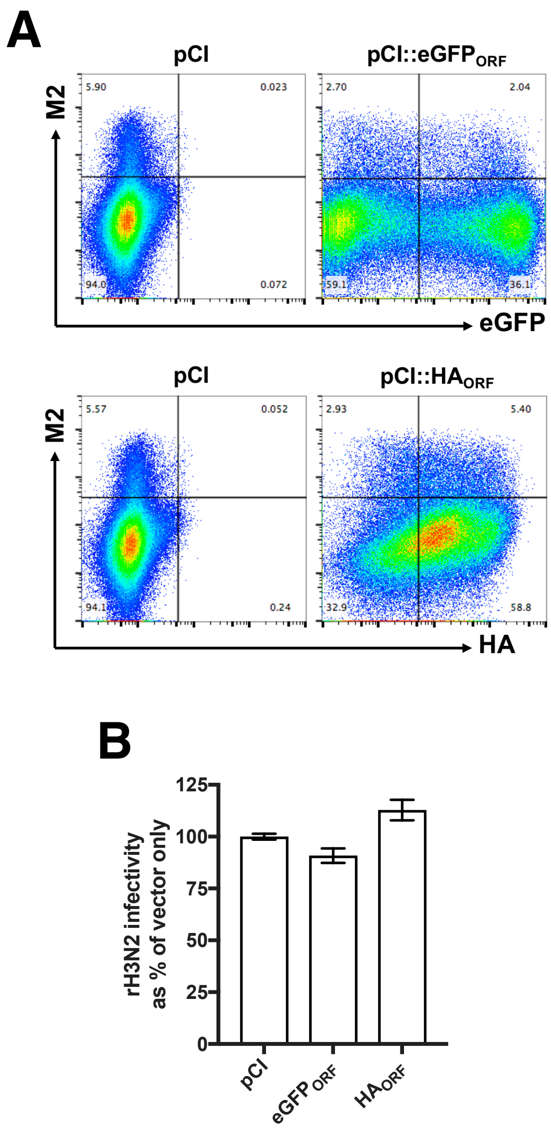


**Figure S5. Overexpression of mRNA and protein does not inhibit subsequent infection. (A)** Cells from the experiment described in Fig 5D are assessed for expression of M2 (indicative of rH3N2 infection) versus eGFP and HA (indicative of transfection) in representative FACS plots. **(B)** rH3N2 infectivity in A are shown as % of pCI vector. Mean values (n=2 cell culture wells) ± standard deviations are shown.
